# The evaluation of RNA-Seq *de novo* assembly by PacBio long read sequencing

**DOI:** 10.1101/735621

**Authors:** Yifan Yang, Michael Gribskov

## Abstract

RNA-Seq *de novo* assembly is an important method to generate transcriptomes for non-model organisms before any downstream analysis. Given many great *de novo* assembly methods developed by now, one critical issue is that there is no consensus on the evaluation of *de novo* assembly methods yet. Therefore, to set up a benchmark for evaluating the quality of *de novo* assemblies is very critical. Addressing this challenge will help us deepen the insights on the properties of different *de novo* assemblers and their evaluation methods, and provide hints on choosing the best assembly sets as transcriptomes of non-model organisms for the further functional analysis. In this article, we generate a “real time” transcriptome using PacBio long reads as a benchmark for evaluating five *de novo* assemblers and two model-based *de novo* assembly evaluation methods. By comparing the *de novo* assmblies generated by RNA-Seq short reads with the “real time” transcriptome from the same biological sample, we find that Trinity is best at the completeness by generating more assemblies than the alternative assemblers, but less continuous and having more misassemblies; Oases is best at the continuity and specificity, but less complete; The performance of SOAPdenovo-Trans, Trans-AByss and IDBA-Tran are in between of five assemblers. For evaluation methods, DETONATE leverages multiple aspects of the assembly set and ranks the assembly set with an average performance as the best, meanwhile the contig score can serve as a good metric to select assemblies with high completeness, specificity, continuity but not sensitive to misassemblies; TransRate contig score is useful for removing misassemblies, yet often the assemblies in the optimal set is too few to be used as a transcriptome.

## Introduction

With the rapid development of sequencing technology, transcriptome assembly by RNA-Seq short reads has become increasingly important in many fields, such as plant science (1; 2), animal science (3) and disease related studies (4; 5; 6). Current transcriptome assembly methods mainly fall into three categories: reference-based assembly, *de novo* assembly and a hybrid assembly that merges the above two (7). For non-model organisms with no available reference genome or transcriptome, *de novo* assembly becomes the only choice to determine the transcriptome before any downstream analysis. Many *de novo* assembly methods have been developed, however, there is no consensus on how to evaluate these methods. Therefore, establishing a reliable benchmark for understanding the property of each *de novo* assembly tool has become a critical issue (8).

Recently, powerful tools, such as Trinity (9), Oases (10), SOAPdenovo-Trans (11), Trans-ABySS (12), and IDBA-Tran (13), have been developed for *de novo* assembly of transcriptomes from RNA-Seq short reads. From the data perspective, when evaluating *de novo* assembly methods, researchers can either simulate RNA-Seq short reads base on a known reference genome or transcriptome (14), or use real RNA-Seq datasets and evaluate the performance of assemblers by comparing the assemblies with the reference transcriptome or the transcriptome of a related species (15; 16). In the first case, even though it is convenient to control the properties of simulated data, such as the expression levels of transcripts, the sequencing error rate, the sequencing depth and etc, the simulated data cannot completely represent the real data. In the latter case, the evaluation heavily relies on the quality of the reference transcriptome. Nevertheless, the expressed transcripts may even vary among biological replicates or different tissues (17). The presence of assemblies that are missed in the reference transcriptome does not necessarily mean that those assemblies are misassemblies. The novel transcript could be representing mutated or fusion transcripts that hasn’t been annotated in the reference. Similarly, the absence of assemblies compared with the reference transcriptome does not necessarily indicate incompleteness of an assembly. It could be that the transcripts that are not expressed in a particular sample. Even though the reference transcriptome is well annotated for a species, e.g., *Homo sapiens*, the reference transcriptomes still vary between different instititional sites (Ensembl and RefSeq) and versions still exist, which complicate the issue from another perspective.

Two types of methods are used for assessing *de novo* assemblies: metrics-based methods and model-based methods. However, without a reference transcriptome, metrics-based methods can only provide an empirical description rather than an assessment of the quality of the assemblies, such as the total number and the length information of assemblies. With a reference transcriptome, the metrics-based methods have the ability to comprehensively evaluate the accuracy, completeness, continuity, and misassembly rate of the assemblies. However, this analysis is based on the assumption that the reference transcriptome is complete and reliable (7; 18). Model-based methods, such as DETONATE (19) and TransRate (20), focus on how well the assemblies can be explained by the read evidence. However, each model-based method has its own definition of the “optimal” assembly, which is inconsistent among different models. Furthermore, model-based methods themselves are hard to evaluate if we do not have a reliable reference transcriptome in hand.

In this study, we utilize the PacBio long read sequencing technology to generate a “real time” transcriptome as a benchmark for assessing (1) the properties of five commonly used *de novo* assembly methods and (2) the effectiveness of two model-based evaluation methods. By comparing the assemblies from the short reads to the “real time” transcriptome from PacBio long reads of the same biological sample, we eliminate the biological uncertainties to a large extent. We conclude that Trinity is best at completeness, but assembled trancripts are less continuous and have more misassemblies than the alternative methods; Oases is best at continuity and specificity (we followed the nomenclature used in rnaQUAST (18); the specificity refers to the percentage of the assemblies that can be well mapped back to the annotated transcripts), but less complete; The performance of SOAPdenovo-Trans, Trans-ABySS and IDBA-Tran are in between. For the model-based evaluation methods, DETONATE ranks the method with all aspects having the average performance as the best, while TransRate doesn’t penalize any downsides but only encourages the good aspects of the assemblies; The contig scores of DETONATE can help select the assemblies with high completeness, specificity and continuity but not a low misassembly rate, while the contig scores of TransRate are helpful in removing misassemblies.

## Methods

### RNA-Seq datasets

The datasets we used were from the Sequencing Quality Control (SEQC)/MAQC-III Consortium, which sequenced a human brain sample by multiple platforms, including MiSeq short read sequencing and PacBio long read sequencing (21). MiSeq generated 7.85 million paired-end reads with the length equal to 250 bp. PacBio generated 0.68 million Reads of Insert (RoIs) with an average length equal to 1, 640 bp.

### Quality control for short read *de novo* assemblies

We first trimmed the adapters and filtered out the low quality reads from the MiSeq dataset using Trimmomatic (version 0.32). Adaptors and low quality reads with average quality below 16 over a 5 base window were removed. And only trimmed reads with length over 30 bases were used for *de novo* assembly. FastQC (version 0.11.2) (https://www.bioinformatics.babraham.ac.uk/projects/fastqc) was then used to visualize the read quality before and after cleaning, shown in S1 Fig.

To determine the best kmer for *de novo* assembly, we used Kmergenie (version 1.6982) (22). Kmergenie examines multiple kmers and counts the frequency of kmers under each k. Then Kmergenie estimates the best k value, which potentially could recover the most possible contigs. Our dataset has the best k = 31 bp, shown in S2 Fig.

Cleaned reads were used for *de novo* assembling by five different assemblers, including Trinity (version 2.2.0), Oases (version 0.2.08), SOAPdenovo-Trans (version 1.03), Trans-ABySS (version 1.5.1), and IDBA-Tran (version 1.1.2). All the methods were tested under the default parameters.

### Quality control for the “real time” transcriptome generated by PacBio long reads

To obtain the real time transcriptome, we ran PacBio long reads through RS_IsoSeq (v2.3.0) pipeline (23) using default parameters. After clustering, we filtered out the non-human genes by aligning both the full length high-quality and full length low-quality consensus sequences to hg19 human reference genome using STAR (24) and GMAP (25) as recommended by RS_JsoSeq. The detailed steps and the number of sequences generated in each step are shown in S3 Fig. Then we collapsed the aligned consenesus sequences by pbtranscript-tofu (https://github.com/PacificBiosciences/cDNA_primer/wiki/tofu-Tutorial-(optional).-Removing-redundant-transcripts) with the minimum alignment identity equal to 0.85 and the minimum coverage to 0.90, as shown in S4 Fig.

## Results

### The real time transcriptome can be served as a reliable benchmark for assessing *de novo* assemblies

Analysis of PacBio long reads with RS_IsoSeq pipeline, produced 9, 636 genes (33, 307 transcripts). All 33, 307 transcripts were corrected versus the hg19 human genome. 244 full length, low quality transcripts can be aligned to hg19 human genome by neither STAR nor GMAP. We pooled these 33, 307 alignable sequences and 244 unalignable sequences together, rendering 33, 551 transcripts and 9, 880 genes in total as the “real time” transcriptome, shown in S4 Fig.

First, to show the relationship between the “real time” transcriptome generated from PacBio long reads and the well annotated human transcriptomes, we drew a venn diagram bewteen the reference transcriptomes from Ensembl and RefSeq and the “real time” transcriptome using vennBLAST (26). Ensembl reference transcriptome has 191, 891 transcripts; RefSeq has 63, 874 transcripts; the “real time” transcriptome has 33, 551 transcripts. In Fig 1, Ensembl has the most transcripts, which almost cover RefSeq and the real time transcriptome. The real time transcriptome is about half the size of RefSeq and largely overlaps with RefSeq. Apparently, the three transcriptomes do not completely overlap each other, which indicates that the evaluations on the *de novo* assemblies would be very different if we chose different reference transcriptomes. Though the “real time” transcriptome is not the most complete set of human transcripts, it derives from the same biological sample as the short reads, which eliminates the uncertainty of sample variance. Therefore, the real time transcriptome should be more optimal as a benchmark for assessing short read assemblies than the other two references.

**Fig 1.**
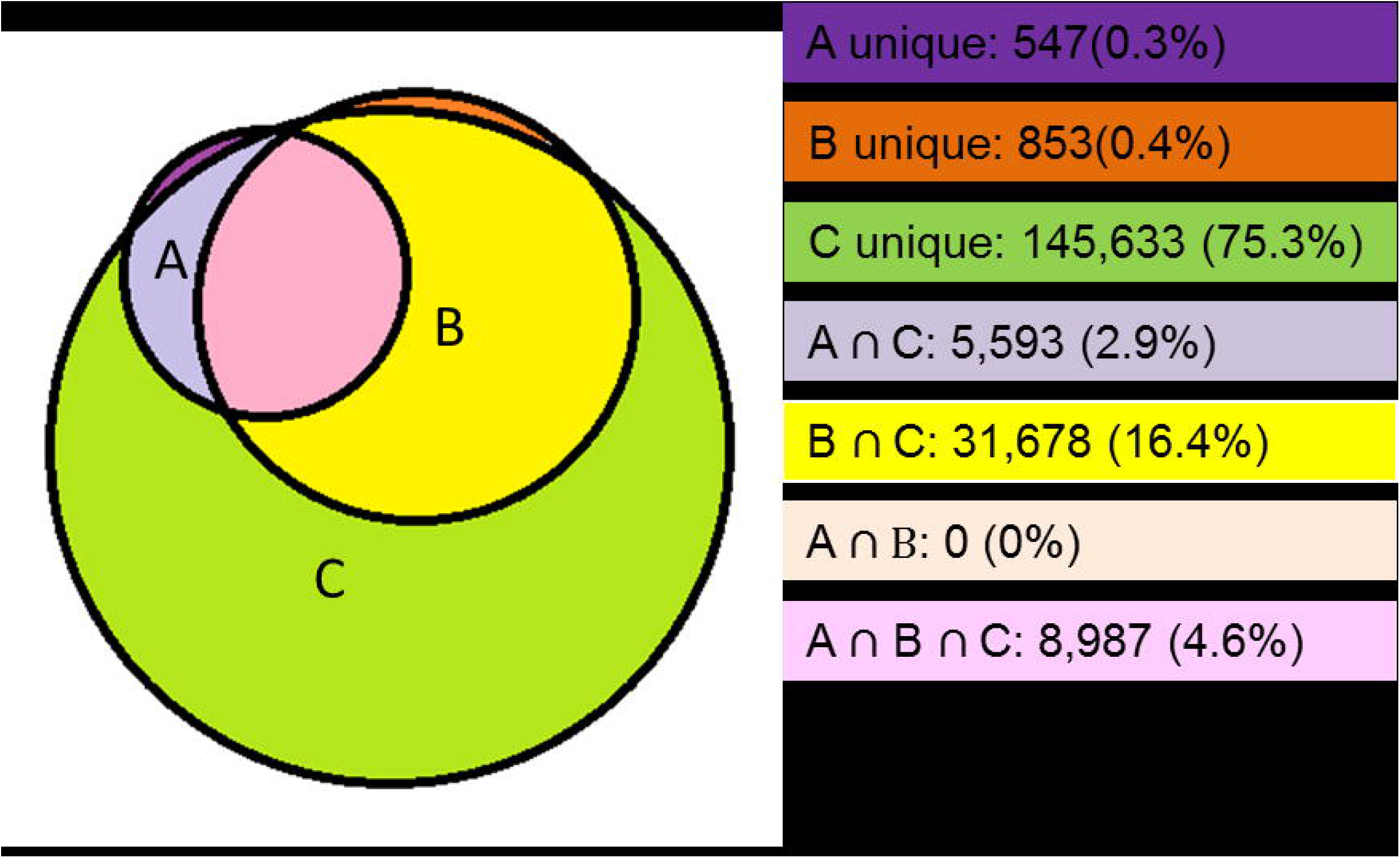
The Venn diagram of three different reference transcriptomes. A is the “real time” transcriptome. B is the RefSeq transcriptome. C is the Ensembl transcriptome. Note that the reason that the total number of transcripts in RefSeq and the real time transcriptome is smaller than the numbers mentioned in the text is because there are multiple transcripts in RefSeq and the real time transcriptome aligning to the same transcript in Ensembl.

Second, we checked whether the abundance of PacBio long reads was corresponds to that of the MiSeq short reads. If yes, it will provide another evidence that the “real time” transcriptome generated from PacBio long reads can serve as a reliable reference for assessing short read assemblies. A scatter plot of the ranks of the abundances estimated by PacBio long reads, and MiSeq short reads is shown in Fig 2. Each data point represents a gene from the “real time” transcriptome. Most highly expressed genes in PacBio also have high expressions as estimated by short reads, and lie in the right up corner. The low expression genes in PacBio have different expression patterns, ranging from low to high as estimated by short reads, and lie along the bottom. This pattern is due to the different throughputs of two sequencing technologies. PacBio has a lower sequencing thoughput than the short read platform. Many transcripts have only one copy detected in PacBio, but these transcripts may have many short reads sampled in MiSeq. This relationship between the abundances of PacBio long reads and MiSeq short reads suggests that a majority of transcripts from the “real time” transcriptome should be recovered by the short read assembly.

**Fig 2.**
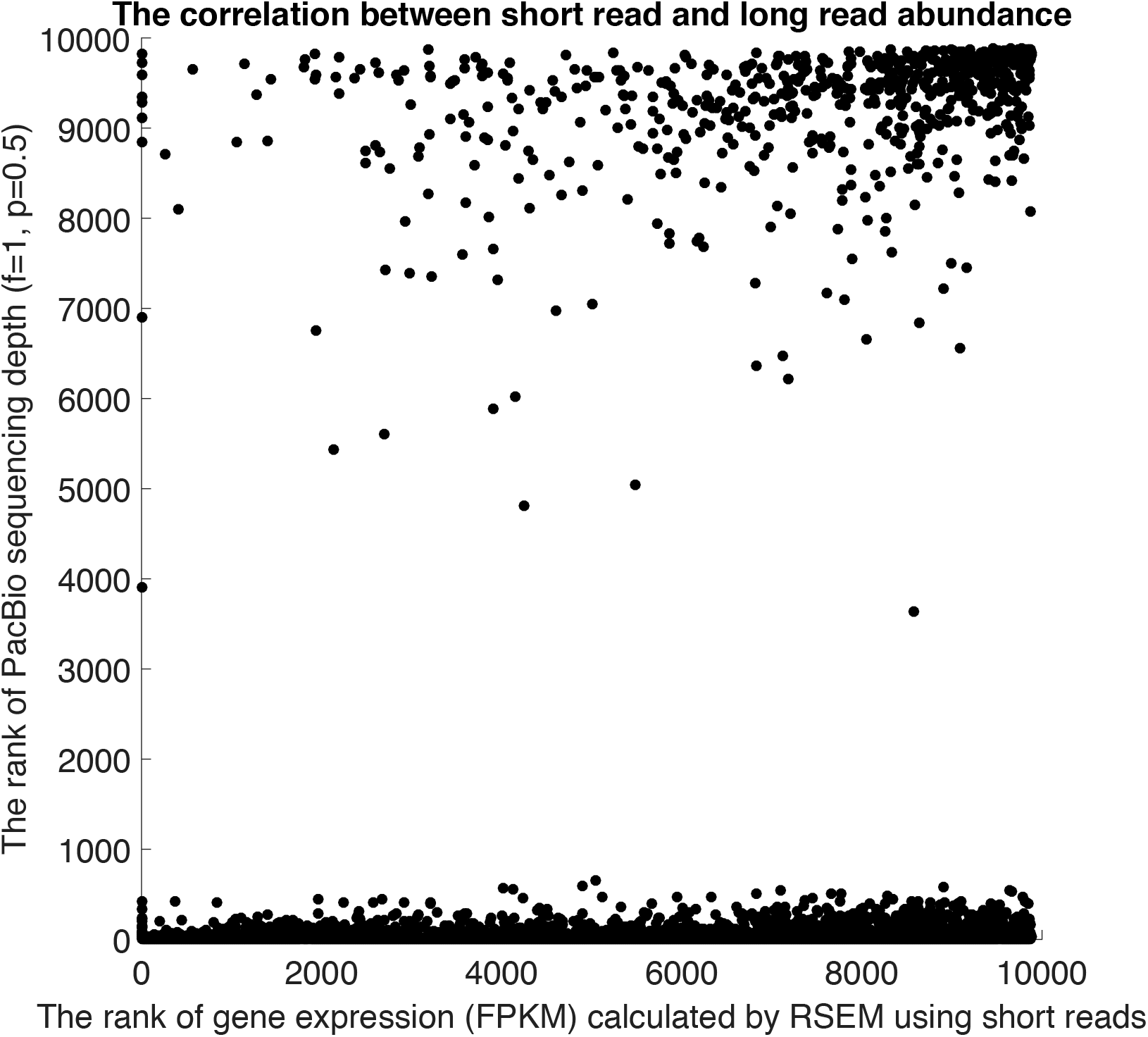
Correlation between the abundance ranks of PacBio long reads and MiSeq short reads. The gene with the lowest abundance is ranked as the first. X-axis shows the ranks of gene expression in Fragments Per Kilobase of transcript per Million mapped reads (FPKM) estimated by RSEM (27) using MiSeq short reads. Y-axis shows the ranks of gene counts from PacBio long reads. If a gene is supported by 1*f*5*p* in PacBio long read sequencing, it means this gene is supported by one full length read and five partial length reads in PacBio. The expression of this gene would be given as (1+ 0.5×5) = 3.5.

In summary, the generation of the real time transcriptome agrees with both the well annotated reference transcriptome and the real time sampling. Therefore, the real time transcriptome can be a better benchmark for assessing short read *de novo* assembling in terms of both sufficency and specificity.

### Assessments on short read *de novo* assembly methods

Assemblies were performed by each method and for each method the number and the length of predicted transcripts are compared. In Table 1, Trinity generates the most assemblies, while Oases generates the fewest assemblies, but with longest average length, median length and N50. The numbers of assemblies in SOAP-denovoTrans, TransAByss and IDBA-Tran are between those of Trinity and Oases. The distribution of the assembly length in Fig 3 shows that the assemblers can be categorized into three groups. Trinity tends to give more assemblies in the range of 200 – 400 bp than alternative methods; Oases tends to give the fewest assemblies in the range of 200 – 400 bp but the curve gradually goes up, having the largest N50 = 1,090 bp. Trans-AByss, SOAPdenovoTrans, and IDBA-Tran share very similar distributions yet IDBA-Tran reports a slightly higher number of assemblies in the range of 300 – 400 bp than Trans-AByss and SOAPdenovoTrans. This finding is consistent with the result in (16), which tested the above assemblers using two authentic RNA-Seq datasets from *Arabidopsis thaliana*. Also, by comparing the assemblies with three reference transcriptomes in Fig 3, including RefSeq, Ensembl, and the “real time” transcriptome, it is clear that all assemblers provide redundant assemblies; and the redundant assemblies are mostly in the short length range.

**Fig 3.**
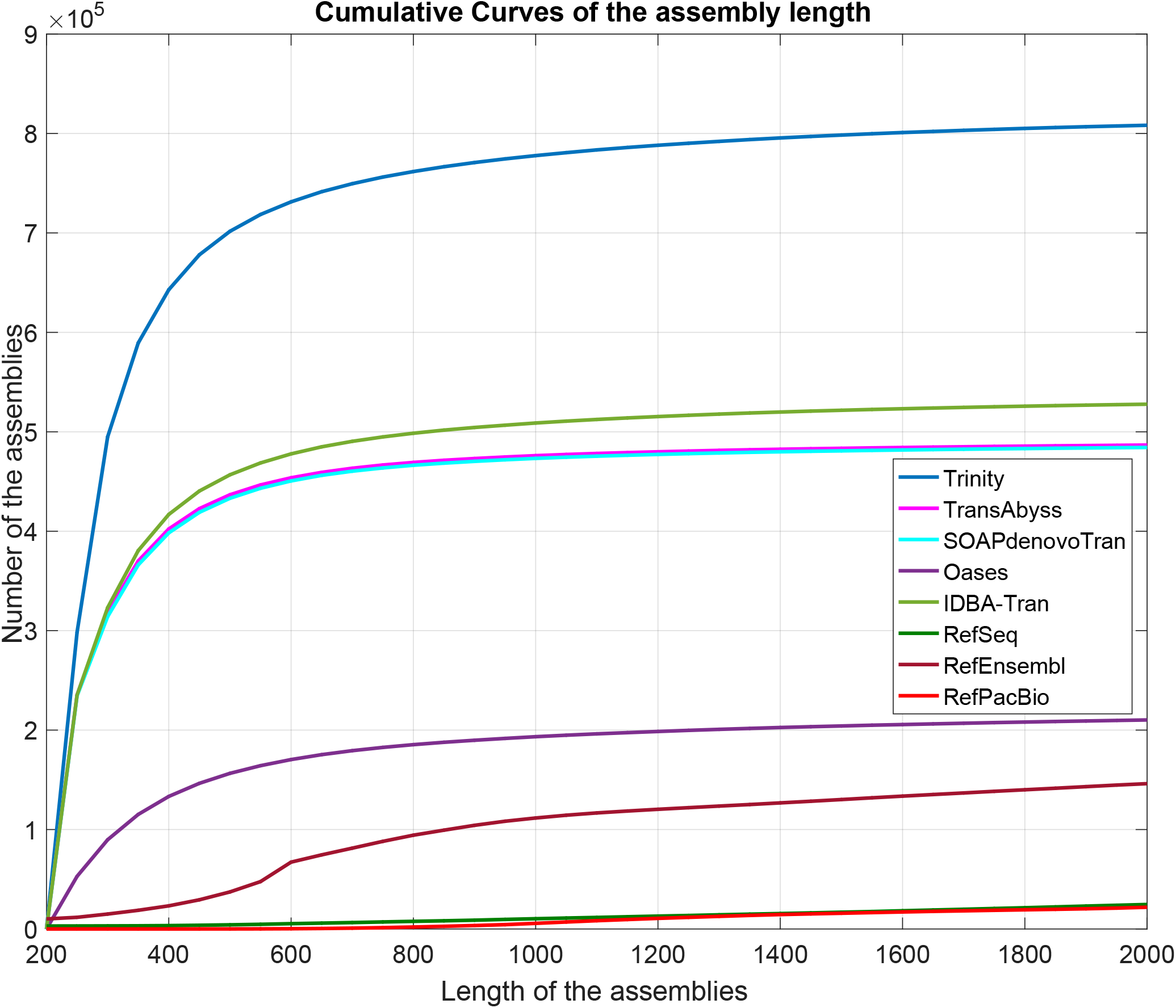
The cumulative curves of the assembly length from five *de novo* assembly methods. There reference transcriptomes are also plotted as quality controls.

**Table 1.** The metrics for the length and the total number of assemblies from five methods.

|  | min_len | max_len | Total bases | The total number of assemblies | avg_len | median_len | N50 |
| --- | --- | --- | --- | --- | --- | --- | --- |
| Trinity | 200 | 22,834 | 332,493,521 | **821,870** | 405 | 273 | 388 |
| IDBA-Tran | 200 | 23,838 | 222,727,826 | 538,261 | 414 | 266 | 412 |
| SOAPdenovo-Trans | 200 | 27,428 | 181,427,977 | 490,473 | 370 | 255 | 350 |
| Trans-AByss | 200 | 27,277 | 322,119,676 | 492,286 | 363 | 254 | 341 |
| Oases | 200 | 27,468 | 149,749,751 | **224,847** | **666** | **343** | **1,090** |

The more short reads that can be aligned back to the assembly, the higher probability that the assembler has generated the correct assembly, if the gene expression level is not taken into account at this stage. Table 2 shows that Trinity, SOAPdenovo-Trans, and Trans-AByss have 75% – 78% short reads that can be mapped back, while Oases and IDBA-Tran only have 56% – 59%. If we only count the number of concordant reads (see the second column in Table 2), the trend is the same as when we count the total number of aligned reads. This suggests that Trinity, SOAPdenovo-Trans and Trans-AByss assemblies potentially contain more information from the short reads than those of Oases and IDBA-Tran. However, if we measure the number of reads mapped back per kilobase of assembly, SOAPdenovo-Trans, Trans-AByss, and Oases have about 60 – 64 short reads mapped back per kilobase of assembly, while IDBA-Tran has 39 and Trinity has only 10. This is because, even though Trinity covers the highest number of short reads, it reports many more predicted assemblies than the other methods.

**Table 2.**
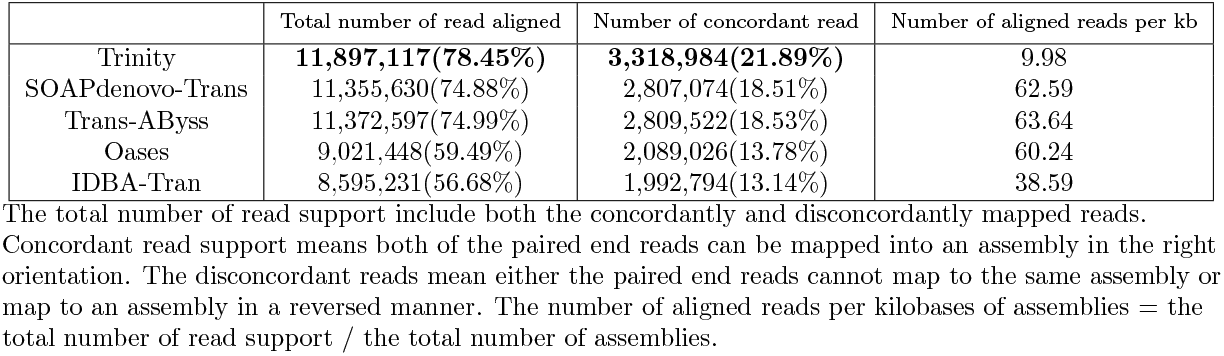
The number of short reads that can be mapped back to the assemblies.

We evaluated the qualities of assemblies by aligning them back to the “real time” transcriptome using rnaQUAST (18) (version 1.4.0). We considered all main statistics reported by rnaQUAST to evaluate the quality of assemblies, including alignability, accuracy, completeness/sensitivity, specificity, continuity, and misassembly. The overall performance of SOAPdenovo-Trans, Trans-AByss and IDBA-Tran (Table 3,) are similar; SOAPdenovo has the highest accuracy and the lowest number of misassemblies in five methods. Oases and Trinity perform very differently, yet each has its own advantages. Oases has the longest average alignment length, the best continuity, specificity, and mean isoform coverage, but Oases assemblies are less complete at both the gene and isoform level. Trinity has the highest completeness at both the gene and isoform level, and a slightly lower specificity than Oases, but a relatively poor continuity and the highest rate of misassemblies. Note that the specificities are very low in all five methods, which indicates a redundancy of assemblies reported.

**Table 3.** The evaluation of the assemblies from five *de novo* assembly methods by comparing with the “real time” transcriptome. We evaluate the assembly quality in terms of the alignability, accuracy, completeness/sensitivity, specificity, continuity, and misassembly. We followed the nomenclatures that were used in rnaQUAST.

|  | SOAPdenovo-Trans | Trans-ABYSS | IDBA-Tran | Oases | Trinity |
| --- | --- | --- | --- | --- | --- |
| **Alignment** |  |  |  |  |  |
| Num. of alignments (hit=50bp) | 489,823(99.87%) | 491,598(99.86%) | 537,458 (99.85%) | 224,564(99.87%) | 820,611(99.85%) |
| Avg aligned length (bp) | 367.8 | 360.1 | 408.8 | 649.8 | 368.4 |
| **Accuracy** |  |  |  |  |  |
| Avg mismatches (bp) per 1K | 1.4 | 1.6 | 1.7 | 1.5 | 3.0 |
| **Completeness/sensitivity** |  |  |  |  |  |
| *Gene level* |  |  |  |  |  |
| Num. of > 50% covered genes | 6,390(66.31%) | 6,385(66.26%) | 6,539(67.86%) | 6,193(64.27%) | 6,895(71.55%) |
| Num. of > 95% covered genes | 2,805(29.11%) | 2,851(29.59%) | 2,796(29.02%) | 2,845(29.52%) | 3,085(32.02%) |
| *Isoform level* |  |  |  |  |  |
| Num. of > 50% covered isoforms | 7,020(21.08%) | 6,952(20.87%) | 7,964(23.91%) | 7,726(23.20%) | 10,396(31.21%) |
| Num. of > 95% covered isoforms | 2,865(8.60%) | 2,885(8.66%) | 2,974(8.93%) | 3,104(9.32%) | 3,423(10.28%) |
| Database coverage | 24.1% | 23.8% | 26.4% | 25.5% | 33.7% |
| Mean isoform coverage | 48.8% | 47.9% | 52.7% | 59.9% | 53.3% |
| **Specificity** |  |  |  |  |  |
| Num. of > 50% matched assemblies | 16,175(3.30%) | 16,309(3.31%) | 24,250(4.51%) | 19,694(8.76%) | 62,758(7.64%) |
| Num. of > 95% matched assemblies | 10,391(2.12%) | 10991(2.23%) | 13003(2.42%) | 9505(4.23%) | 35316(4.30%) |
| Mean fraction of assemblies matched | 3.4% | 3.4% | 4.5% | 8.6% | 7.6% |
| Unannotated assemblies | 459,792(93.74%) | 458,408 (93.12%) | 492,515(91.50%) | 192,489(85.61%) | 708,017 (86.14%) |
| **Continuity** |  |  |  |  |  |
| *Gene Level* |  |  |  |  |  |
| Num. of > 50% assembled genes | 5,228(54.25%) | 5,243(54.41%) | 5,266(54.65%) | 5,370(55.73%) | 4,967(51.55%) |
| Num. of > 95% assembled genes | 2,270(23.56%) | 2,243(23.28%) | 2,123(22.03%) | 2,381(24.71%) | 1,800(18.68%) |
| *Isoform Level* |  |  |  |  |  |
| Num. of > 50% assembled isoforms | 5,604(16.83%) | 5,561(16.70%) | 6,287(18.88%) | 6,663(20.00%) | 7,160(21.50%) |
| Num. of > 95% assembled isoforms | 2,313(6.94%) | 2,268(6.81%) | 2,265(6.80%) | 2,605(7.82%) | 2,002(6.01%) |
| Mean isoform continuity | 42.4% | 41.5% | 45.5% | 54.3% | 42.9% |
| **Misassemblies** | 3,233(0.66%) | 5,859(1.19%) | 9,874(1.83%) | 4,332(1.93%) | 30,915(3.76%) |

### Assessments of model-based *de novo* assembly evaluation methods

There are two state-of-the-art methods having been assessed here, DETONATE (version 1.9) and TransRate (version 1.0.1). The RSEM-EVAL module of DETONATE is used for evaluating *de novo* assemblies without a reference transcriptome. The RSEM-EVAL score is the sum of three components; the likelihood estimates how well the assemblies are explained by the mapped short reads; the assembly prior assumes the assembly length follows a negative binomial distribution and the transcripts are independent from each other (the number of isoforms or homogenous genes will influence this component); the BIC penalty penalizes the prediction of too many bases and assemblies. Table 4 shows that the likelihood makes the largest contribution to the RSEM-EVAL score. Consistent with Table 2 and 3, Trinity has the highest likelihood, but the lowest assembly prior and BIC penalty, which lowers its overall RSEM-EVAL score. On the contrary, SOAPdenovo-Trans and Trans-AByss do not score highly any component, which is consistent with Table 3, but achieve the best overall RSEM-EVAL score, because no single component dominates the final evaluation. IDBA-Tran and Oases have low RSEM-EVAL scores mainly due to the low likelihoods, though Oases has the best assembly prior and BIC penalty, which is also consistant with Table 2 and 3.

**Table 4.** DETONATE RSEM-EVAL scores for five *de novo* assembly methods.

|  | SOAPdenovo-Trans | Trans-ABYSS | IDBA-Tran | Oases | Trinity |
| --- | --- | --- | --- | --- | --- |
| Likelihood | -3,001,246,561 | -3,016,249,561 | -3,238,791,356 | -3,225,800,940 | **-2,841,562,473** |
| Assembly prior | -251,577,741 | -247,841,006 | -308,900,873 | **-207,358,450** | -461,191,586 |
| BIC penalty | -3,884,882 | -3,899,242 | -4,263,395 | **-1,780,946** | -6,509,768 |
| RSEM-EVAL score | **-3,256,709,184** | -3,267,989,809 | -3,551,955,624 | -3,434,940,336 | -3,309,263,827 |

The RSEM-EVAL score is the sum of three components, including the likelihood, the assembly prior and the BIC penalty for each assembly set.

TransRate shows the opposite pattern compared with DETONATE. The TransRate assmbly score is the geometric mean of the contig scores multiplied by the proportion of short reads that positively support the assemblies. Each contig score is the product of four components: the nucleotide score measuring the alignment distance between the assembly and the short reads, the coverage score measuring the fraction of the assembly length covered by reads, the order score measuring the orientation of the paired-end read mapping, and the segment score measuring the per-nucleotide read coverage. In Table 5, we find that Oases has the highest TransRate score. However, after optimizing an empircal target function – 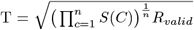, where *S*(*C*) is the contig score, *n* is the number of selected contigs, and *R_valid_* is the proportion of reads that can be mapped to the selected contigs – as recommended by TransRate, the TransRate scores of all methods greatly increase, and Trinity has the best optimal score.

**Table 5.**
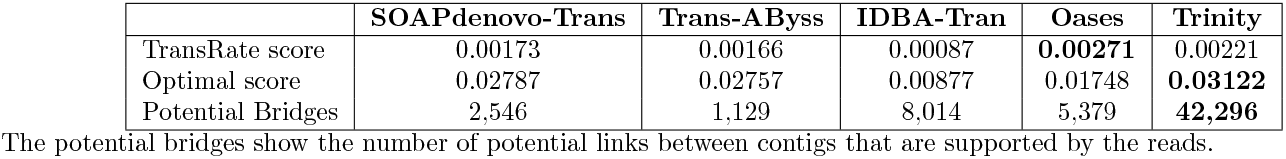
TransRate assembly and the assembly scores after optimization for five *de novo* assembly methods.

### Contig scores can serve as a good metric for removing low quality assemblies

In the section of assessments on short read *de novo* assembly methods, we found that the number of predicted assemblies produced by *de novo* approaches was about 15 times of the number of transcripts in the”real time” transcriptome, on average; In the section of assessments of model-based *de novo* assembly evaluation methods, we found that a large portion of assemblies have low DETONATE and TranRate contig scores. Together, this indicates a redundency of assemblies. Therefore, the question is whether DETONATE and TransRate contig scores can serve as good metrics for removing low quality assemblies.

We selected the top 40, 000 assemblies based on the DETONATE score, the TransRate score, and the FPKM of each assembly. An ideal selection would be an assembly set with no change in completeness, but with increased specificity and continuity, and decreased misassembly rate, compared with the full set of assemblies. Fig 4 shows the comparison between the full set and the selected set of assemblies in the completeness, specificity, continuity and the misassembly rate. For completeness, the database coverage rates are decreased in all selected sets compared to the full set, but DETONATE selections show higher completeness than TransRate and FPKM. For specificity, DETONATE selections show a generally higher mean fraction of matched assemblies than the full set of assemblies in all methods, but not the other two metrics. For continuity, DETONATE selections also show a generally higher mean fraction of isoform length assemblied than the full set of assemblies; FPKM selections also have a higher continuity than the full set in all methods, except for Trinity. For the misassembly rate, both TransRate and FPKM selections can greatly decrease the number of misassemblies but not DETONATE, which might because TransRate takes the order score into account.

**Fig 4.**
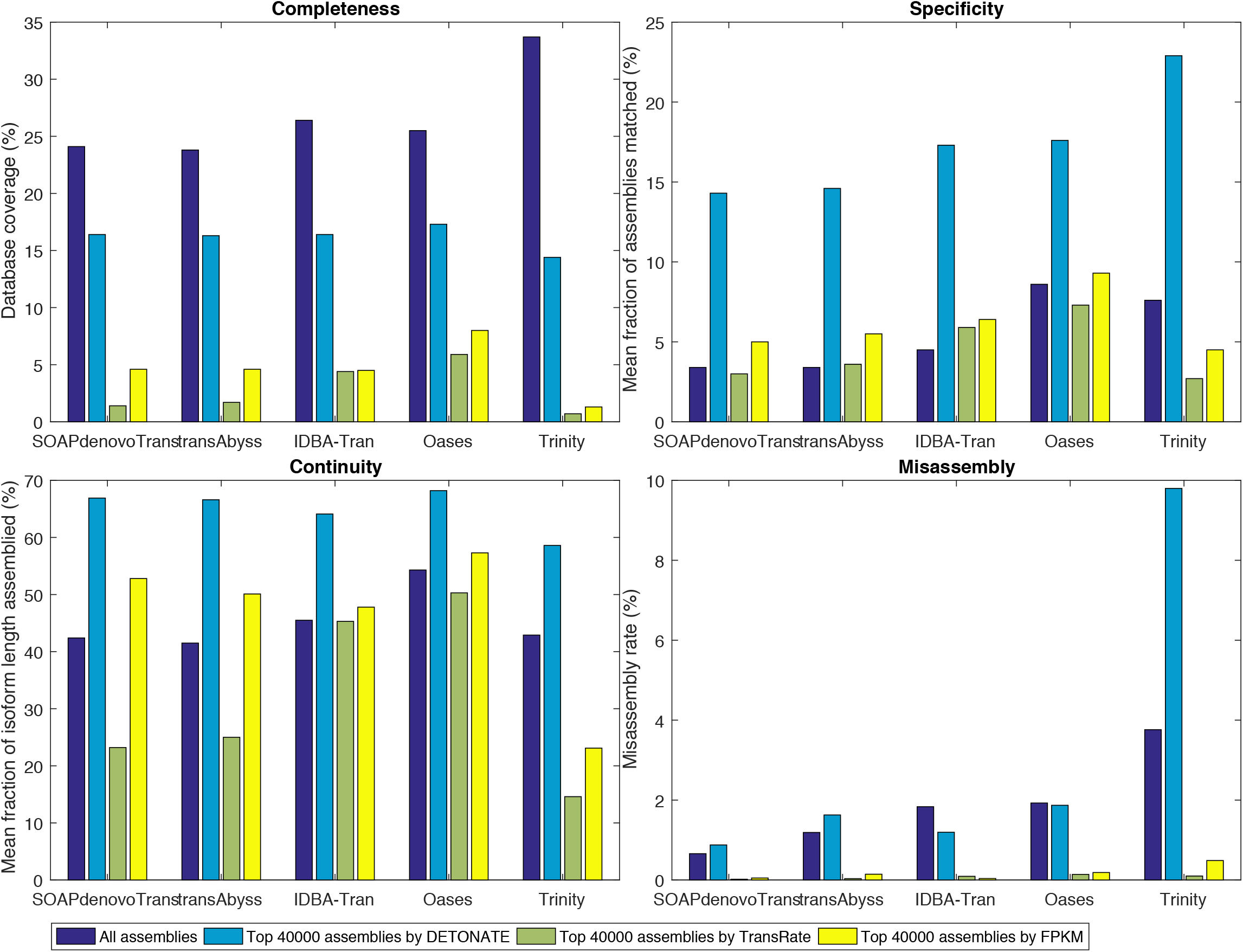
The efficiency to remove low quality assemblies by three different metrics, including DETONATE contig score, TransRate contig scores and the FPKMs of contigs. Four major aspects have been evaluated by comparing the top 40, 000 selected assemblies with the”real time” transcriptome. We evaluate the assembly quality in terms of the completeness, specificity, continuity, and the misassembly rate, as that in Table 3. The misassembly rate is calculated as the number of misassemblies divided by the number of assemblies.

## Discussion

In this study, we propose a reliable benchmark – a real time transcriptome, produced by PacBio long read sequencing – for assessing the *de novo* assembly and evaluation methods. As opposed to other *de novo* assembly assessment strategies, which either simulate RNA-Seq data or utilize well annotated reference transcriptome as a groud truth for real data, our study takes the advantage of sequencing the same biological sample using both the short read and long read technologies to eliminate the biological uncertainty. The real time transcriptome relies on both the well annotated reference transcriptome and the real time sampling, thus, the real time transcriptome can serve as a better reference for assessing *de novo* assemblies than the alternative simulation or a reference transcriptome.

By comparing the *de novo* assemblies from five commonly used methods to the real time transcriptome, we find that the properties of the tested assemblers vary significantly. For instance, Trinity has the highest read mapping rate (shown in Table 2), and the best completeness, but generates too many short assemblies in the range between 200 – 400 bases (shown in Fig 3). This makes Trinity assemblies less continuous, and potentially increasing the number of assemblies that can be linked by short reads (shown in Table 4). Trinity also has the highest misassembly rate of the five methods (shown in Table 3). An improvement to Trinity would be to decrease the number of misassemblies while increasing the continuity. Oases generally generates the longest and the fewest assemblies in all five methods (shown in Table 1), which gives it the best continuity and specificity (shown in Table 3). However, Oases has a low read mapping rate (shown in Table 2), which makes it less complete than the other methods. An improvement to Oases would be to increase the completeness of the assemblies. The performance of SOAPdenovo-Trans, Tran-AByss, and IDBA-Tran are very similar, but SOAPdenovo-Trans has the lowest number of mismatches and misassemblies of the five methods (shown in Table 3).

Because of the overall redundency of *de novo* assemblies in all the methods, DETONATE and TransRate can serve as good metrics to evaluate and remove low quality assemblies, but with different patterns. The DETONATE assembly score mainly considers the read mapping rate, the independence of trancripts, the total number of assemblies, and the number of assembled bases when evaluates the assemblies. DETONATE ranks the method with no extreme disadvantages in the above aspects as the best (shown in Table 4). The TransRate assembly score is an empirical function that takes many different aspects into account, mainly including the mapping accuracy, the mapping orientation, the mapping depth, the mapping coverage, and the fraction of mapped reads. By taking the product of the first four terms as the contig score, TransRate actually treats the first four aspects equally, then weights the contig score by the fraction of mapped reads. TransRate only encourages the advantages but doesn’t penalize the disadvantages of the assembly, as the way DETONATE does. The optimization of the TransRate assembly score is a good way to select the best quality assemblies, but the number of selected assemblies is often low, and cannot be controlled by users.

Both DETONATE and TransRate provide contig scores as an evaluation for each assembly. The contig scores can be used as metrics for removing redundent low quality assemblies. When the top 40, 000 assemblies ranked by DETONATE, TransRate and FPKM are examined, we find that the DETONATE contig score can effectively remove the redundent assemblies while keeping a high completeness and continuity rate, but not be able to remove misassemblies. The TransRate contig score is very sensitive in removing misassemblies but not helpful in the completeness, specificity and continuity.

There are some weakness in this study. For instance, only one dataset has been tested here, because it is not very easy to obtain the datasets which have been sequenced by both short read and long read technologies. It would be better to include further benchmark datasets to eliminate any bias from the sequencing platforms or organisms. Also, we evaluated the assemblies from several major perspectives, including length, the total number of assemblies, the read mapping rate, completeness, specificity, continuity and misassembly, by comparing the assemblies with the real time transcriptome. There may be additional perspectives that the model-based evaluation methods take into account, but are not included in our metrics.

## Supporting information

**S1 Fig. MiSeq read quality visualization by FastQC before and after trimming.** (A) and (B) are the positional qualities of forward and backward reads in the raw dataset. (C) and (D) are the positional read qualities of forward and backward reads after trimming.

**S2 Fig. Kmergenie shows kmer = 31bp is the best choice for short read assembly.**

**S3 Fig. The flowchart of processing PacBio long reads into real time transcriptome.** We begin from the .bax.h5 raw data. The first step is classification, namely to classify Reads of Inserts (RoIs) into full-length and non-full-length RoIs based on the adaptors, meanwhile removing the chimeric RoIs. The second step is clustering, namely to cluster RoIs into consensus, while each consensus can be viewed as a transcript. The third step is collapsing and correction, namely to align the consensus sequences back to the reference genome and get the real time transcriptome.

**S4 Fig. The selection of the thresholds for coverage and identity when align the consensus sequences back to the human reference genome.** Because the coverage and identity are the only parameters the user has to set when running the collapse step in pbtranscript-tofu.py, and these parameters will eventually influence the numbers of transcripts and genes in the real time transcriptome, we carefully select the values of these two parameters. (A) and (B) show the number of transcripts and genes in the real time transcriptome when we set different coverages and identities, respectively. Note that when the coverage is from 0.90 to 0.95, more transcripts/genes will be dropped out than the previous columns, which indicates many consensus sequences having a coverage between 0.90 and 0.95. To keep as much information from PacBio long reads as possible, we consider coverage = 0.9 is a long enough to represent a transcript/gene. Similarly for identity, when identity is from 0.85 to 0.90, more transcripts/genes will be dropped out than the previous columns. Taking the facts that the error rate of PacBio sequencing was about 10%-15% in 2013 and the average length of consensus sequence (1, 640 bp) was long enough to align the consensus sequence to the right position on the genome into account, we choose identity = 0.85 in our dataset.

## Acknowledgments

We acknowledge Dr. Christopher Mason in Cornell University and Dr. Farmerie William in the University of Florida for providing the raw datasets of PacBio long read sequencing.

