## Supplementary figures and images for "The evaluation of RNA-Seq *de novo* assembly by PacBio long read sequencing"

### S1.Fig

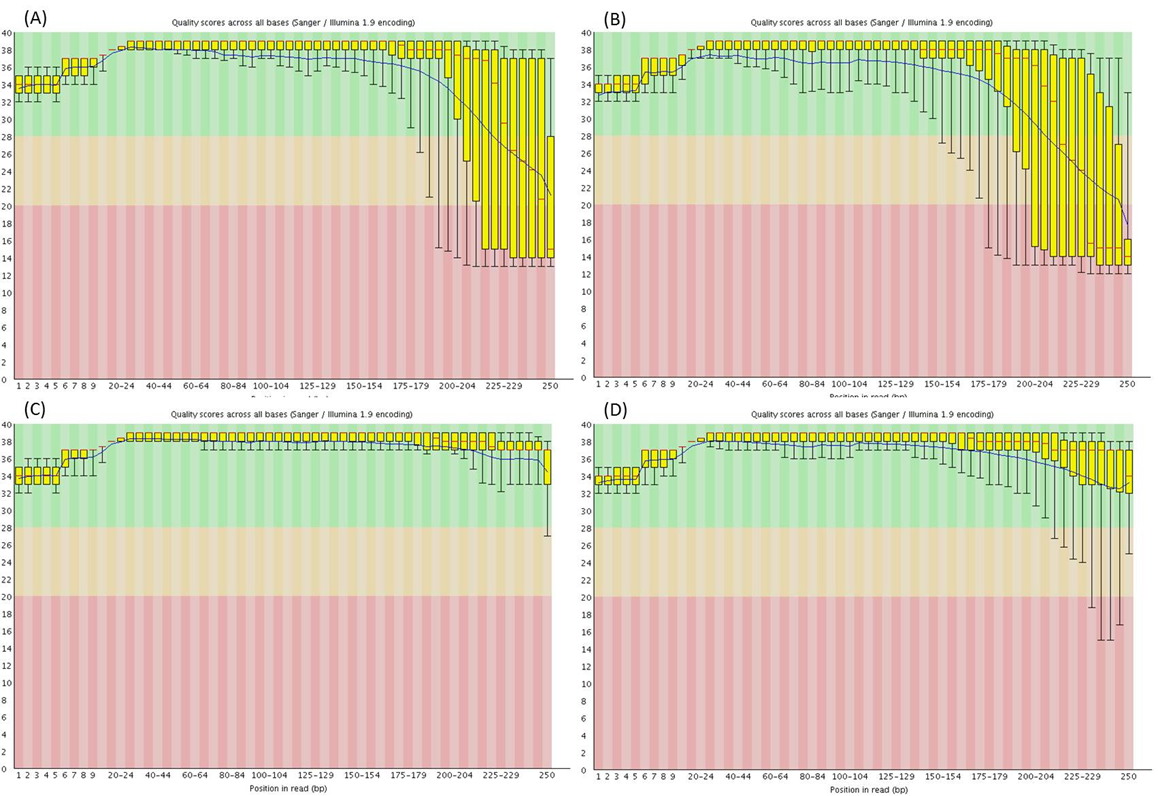

### S2.Fig

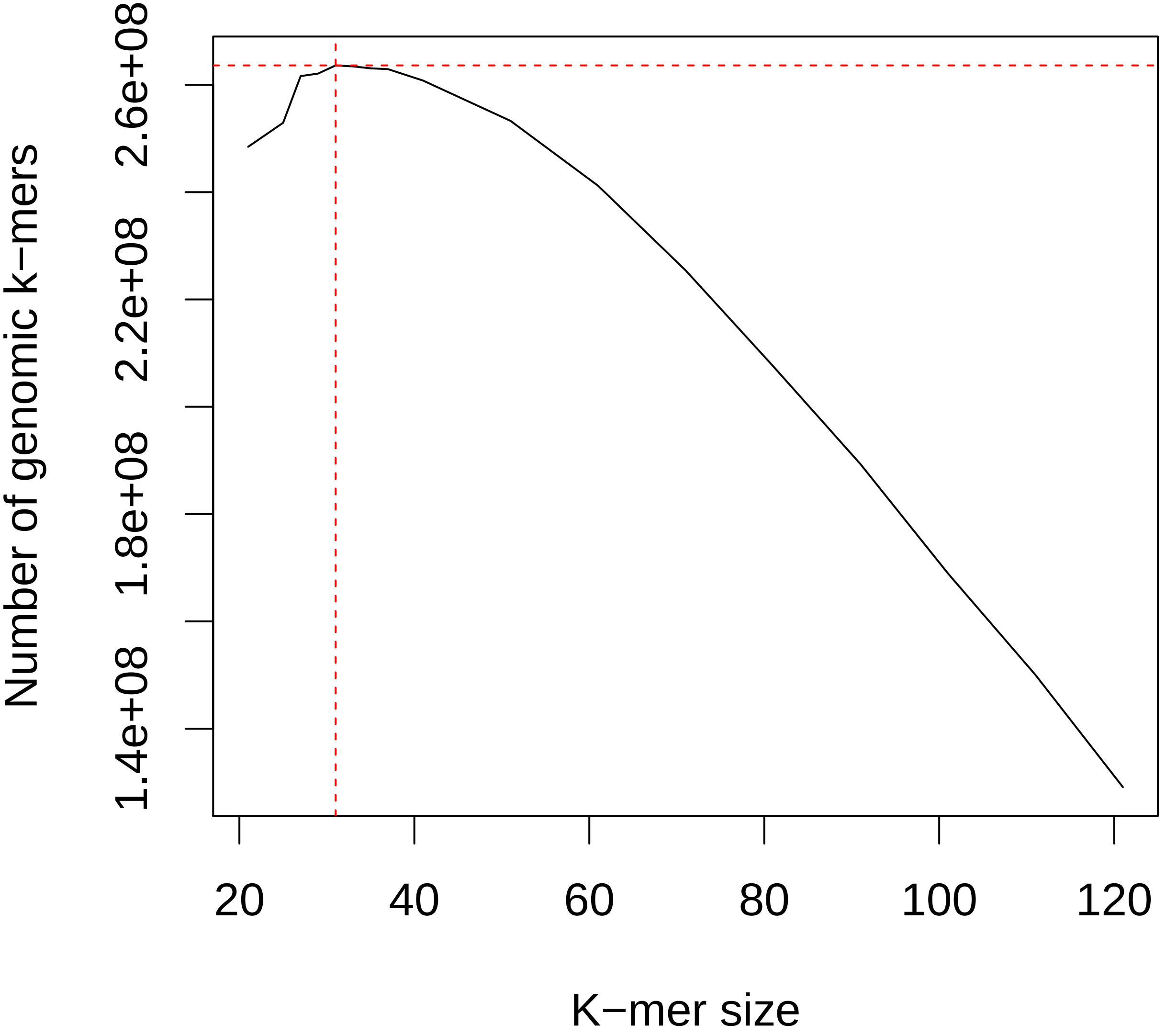

### S3.Fig

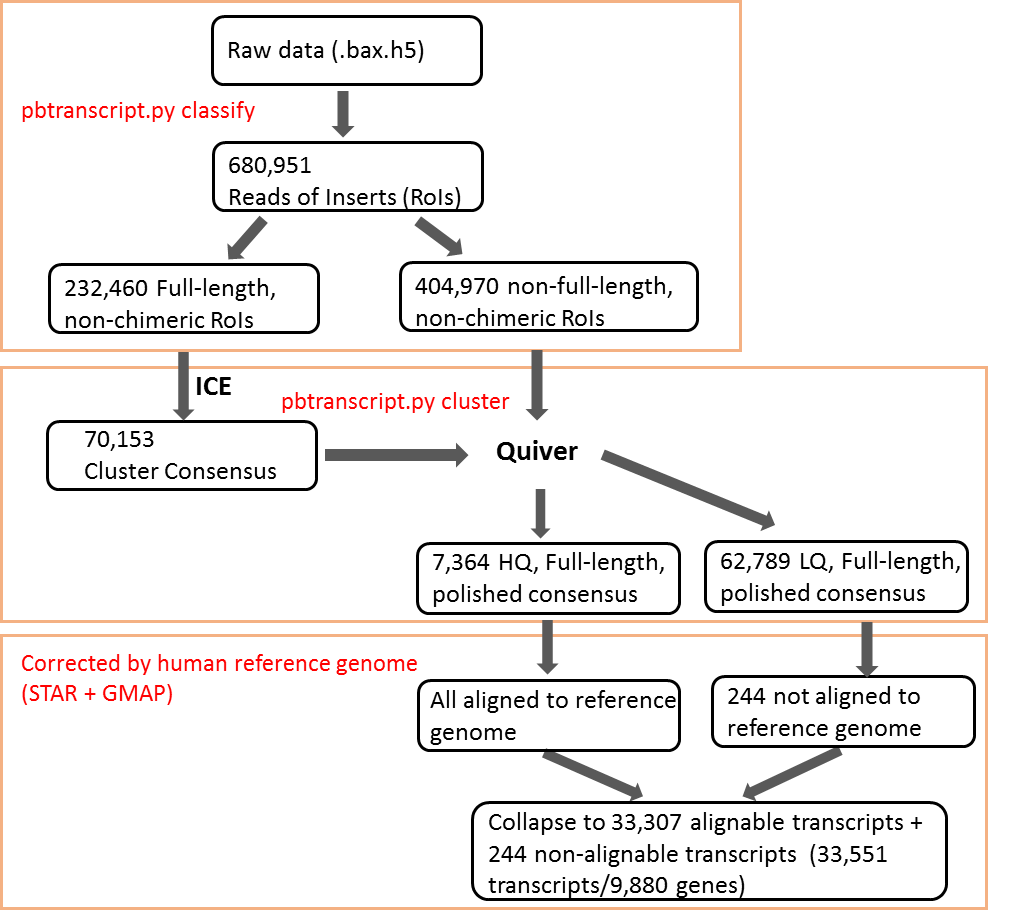

### S4.Fig

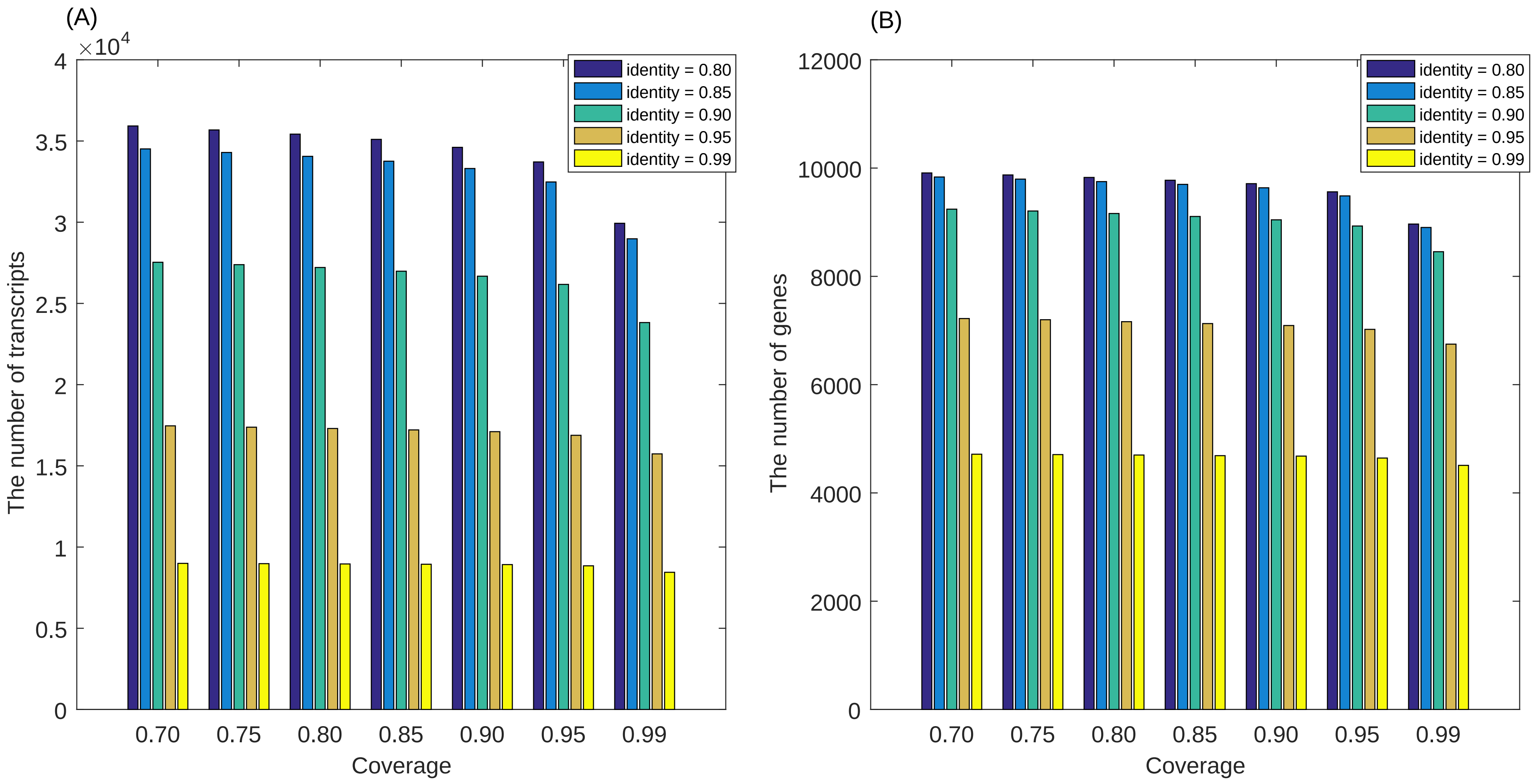
